# Predicting time to dementia using a quantitative template of disease progression

**DOI:** 10.1101/458273

**Authors:** Bilgel Murat, Bruno M. Jedynak

## Abstract

**Introduction:** Characterization of longitudinal trajectories of biomarkers implicated in sporadic Alzheimer’s disease (AD) in decades prior to clinical diagnosis is important for disease prevention and monitoring.

**Methods:** We used a multivariate Bayesian model to temporally align 1369 AD Neuroimaging Initiative participants based on the similarity of their longitudinal biomarker measures and estimated a quantitative template of the temporal evolution cerebrospinal fluid (CSF) A*β*_1-42_, p-tau_181p_, and t-tau, hippocampal volume, brain glucose metabolism, and cognitive measurements. We computed biomarker trajectories as a function of time to AD dementia, and predicted AD dementia onset age in a disjoint sample.

**Results:** Quantitative template showed early changes in verbal memory, CSF A*β*_1-42_ and p-tau_181p_, and hippocampal volume. Mean error in predicted AD dementia onset age was < 1.5 years.

**Discussion:** Our method provides a quantitative approach for characterizing the natural history of AD starting at preclinical stages despite the lack of individual-level longitudinal data spanning the entire disease timeline.

## 1. Background

Alzheimer’s disease (AD) related brain changes, including amyloid and phosphorylated tau deposition as well as neurodegeneration, begin years prior to the emergence of clinical dementia [1]. There is great interest in understanding the progression of biological and cognitive markers (collectively referred to as biomarkers in this paper) that are implicated in AD in the earliest stages, given that therapeutic intervention is hypothesized to be more effective if administered early in the disease before downstream brain damage occurs. Such an understanding will enable a better definition of preclinical AD and help identify individuals who are likely to benefit from therapy.

In studies of dominantly inherited AD, parents’ ages at disease onset serve as strong predictors of an individual’s onset age, allowing the use of time from expected onset as a surrogate of disease progression against which biomarker trajectories can be characterized [2]. However, in sporadic AD, such an estimate is not as good an indicator of expected onset, making it difficult to characterize biomarker evolution in the earliest stages of neuropathology and neurodegeneration that mark the preclinical stages of AD. Furthermore, differences across individuals in the rate of disease progression make it difficult to characterize a quantitative template (QT) of biomarker changes as a function of disease state. Therefore, it is necessary to use novel statistical approaches that take into account such differences across individuals for combining short-term follow-up data per individual to reveal long-term biomarker trajectories.

Several studies have addressed this challenge of characterizing biomarker trajectories in preclinical stages of AD as a function of an underlying latent disease progression variable that reflects the natural history of AD neuropathology, neurodegeneration, and cognitive changes via generative models. These models emphasize the interpretability of model results rather than optimization of predictive performance, where discriminative methods might outperform but produce results that are not as easily interpretable. Existing generative models of AD progression can be divided into two major categories based on the granularity of their characterization of the latent disease progression variable, either as a sequence of discrete events [3–5] or as a continuous variable [6–14]. In addition to characterizing changes in biomarker trajectories as a function of latent disease stages, these statistical models provide individualized information that can be used for personalized disease staging and monitoring.

Recent work in this area has focused on Bayesian reformulations of statistical approaches to biomarker trajectory modeling [11, 14, 15] to enable probabilistic estimates of trajectories and better characterization of the individual-level uncertainty in disease progression variables. These improvements can lead to better disease monitoring and progression prediction at the individual level, thereby providing useful tools for clinical trial recruitment and assessment. We further this line of research into continuous latent disease progression models by reformulating our Progression Score Model [6, 7] as a Bayesian model, and make substantial changes to improve the interpretability of our results. First, we impose weakly informative priors on model parameters. This allows us to compute credible intervals around estimated trajectories and individualized disease stage indicators. Second, based on our earlier observation that rate of progression is associated with disease stage, we revise the transformation between age and the latent disease progression variable, so that biomarker trajectories can be depicted as a function of time to diagnosis, which was not directly possible in our previous model.

Using this Bayesian Progression Score Model, we compute a QT of the temporal evolution of AD-related biomarkers by temporally aligning longitudinal data for individuals based on the similarity of a collection of biomarkers for individuals who are cognitively normal, have mild cognitive impairment (MCI) or AD dementia. The QT reflects change in biomarkers as a function of a latent disease stage indicator that is simultaneously estimated per individual in the model. Unlike Bayesian models proposed by Li et al. [11] or Lorenzi et al. [15], where estimated biomarker trajectories are presented only in terms of the latent disease progression variable, our model enables the characterization of the trajectories in time domain, thereby facilitating the interpretation and applicability of our results to clinical settings. Our use of basis functions for depicting the time courses of biomarkers allow to capture more flexible trajectories than those possible with the parametric form proposed by Ishida et al. [14]. The estimated temporal QT shows that verbal memory decline is detectable early on the trajectory to MCI and AD. The temporal evolution is also marked by changes in hippocampal volume, cerebrospinal fluid (CSF) amyloid (A*β*_1-42_), total tau (t-tau) and phosphorylated tau (p-tau_181p_), as well as brain glucose metabolism and global measures of cognition and mental status. We demonstrate that the estimated latent disease progression variable is associated with known risk factors for late-onset AD and clinical diagnosis. Finally, we predict AD dementia onset age using baseline data in a disjoint testing set, achieving a root mean square error of < 1.5 years. Our results provide insights into the natural history of late-onset Alzheimer’s disease starting with the preclinical stages. The estimated QT can be used to estimate individualized latent disease stages given a collection of biomarker measurements and predict future conversion to AD.

## 2. Method

Data used in the preparation of this article were obtained from the Alzheimer’s Disease Neuroimaging Initiative (ADNI) database (adni.loni.usc.edu). The ADNI was launched in 2003 as a public-private partnership, led by Principal Investigator Michael W. Weiner, MD. The primary goal of ADNI has been to test whether serial magnetic resonance imaging (MRI), positron emission tomography (PET), other biological markers, and clinical and neuropsychological assessment can be combined to measure the progression of MCI and early AD. Specifically, we used the ADNI data prepared for the Alzheimer’s Disease Modelling Challenge and followed the recommendations prescribed in this document for easing the comparison with other quantitative templates for the progression of AD.

### Participants

We used data for 1706 participants with 6880 visits from ADNI1/GO/2. Clinical diagnoses of MCI and AD dementia were determined by the ADNI Conversion Committee according to the criteria described in the ADNI protocol. We designated data for approximately 80% of the individuals selected at random as the training set (1369 individuals with 5533 visits). We designated the data for the remaining individuals, excluding 67 individuals with AD at baseline, as the testing set (270 individuals with 1165 visits) for evaluating the performance of age at AD onset prediction. Participant demographics are presented in Table 1. We compared demographic variables by diagnostic category using Wilcoxon rank-sum test for continuous variables and Fisher’s exact test for categorical variables.

**Table 1:** Participant demographics at baseline. Continuous variables are reported as “mean (standard deviation)” and categorical variables are reported as “count (percentage)”.

|  | *Training set* |  |  | *Testing set* |  |
| --- | --- | --- | --- | --- | --- |
|  | NL<br>*n* = 405 | MCI<br>*n* = 698 | AD<br>*n* = 266 | NL<br>*n* = 114 | MCI<br>*n* = 156 |
| Age | 74.5 (5.81) | 72.8 (7.62) | 74.9 (7.71) | 73.3 (5.78) | 73.5 (7.40) |
| Male | 197 (48.6%) | 413 (59.2%) | 149 (56.0%) | 56 (49.1%) | 91 (58.3%) |
| White | 372 (91.9%) | 651 (93.3%) | 246 (92.5%) | 100 (87.7%) | 146 (93.6%) |
| Education | 16.3 (2.62) | 15.9 (2.85) | 15.3 (3.02) | 16.5 (2.97) | 16.0 (2.85) |
| Single $\epsilon 4$ | 105 (26.0%) | 280 (40.2%) | 132 (50.0%) | 32 (28.1%) | 56 (35.9%) |
| $\epsilon 4/\epsilon 4$ | 7 (1.73%) | 75 (10.8%) | 46 (17.4%) | 5 (4.39%) | 18 (11.5%) |
| No. visits | 4.21 (2.28) | 4.45 (1.97) | 2.71 (1.14) | 4.25 (2.27) | 4.37 (2.04) |
| Follow-up | 3.15 (2.38) | 2.75 (1.84) | 1.14 (0.84) | 3.14 (2.44) | 2.80 (1.99) |

### Biomarkers and cognitive measures

We computed a QT of the temporal evolution of the following AD-related biomarkers: CSF A*β*_1-42_, p-tau_181p_, t-tau; intracranial volume-adjusted hippocampal volume; brain glucose metabolism measured by fluorodeoxyglucose (FDG) PET; verbal memory measured by Rey Auditory Verbal Learning Test (RAVLT) immediate recall (sum score across 5 learning trials) [16]; mental status measured by the Mini Mental State Exam (MMSE) [17]; and AD and dementia indicators as measured by the Alzheimer’s Disease Assessment Scale-Cognitive 13-item scale (ADAS13) [18], and the Clinical Dementia Rating-Sum of Boxes (CDR-SB) [19]. These measures were selected based on their demonstrated involvement in the progression of AD, and closely mirror those selected for analysis with an earlier version of our model [6]. We selected visits with at least 5 of these 9 measures for inclusion in our analysis.

### Progression score model

We reformulated the previously described Progression Score Model [6, 7] using a Bayesian framework where biomarker trajectories are modeled using basis functions. Compared to linear or sigmoid functions used in our previous applications, basis functions allow for much richer models of biomarker trajectories. We make the following modeling assumptions:

1. Changes in biomarkers relative to one another in the progression from a cognitively normal state to AD dementia can be characterized uniquely.
2. Any deviation from this unique characterization of biomarker changes along the disease spectrum is assumed to be measurement noise.
3. Given enough time, all individuals will exhibit biomarker levels seen in AD dementia. However, an individual’s progression might be slow enough that AD-level biomarkers will not be attained during the individual’s life span.

### Continuous-time model

We first describe the model in continuous time. We then discretize it for fitting the parameters. The model describes biomarker values collectively as a function of age using a two-level composition. First, there is a subject-dependent exponential mapping of age *t* to progression score (PS) *s*:

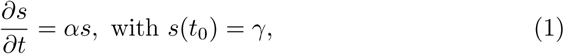

where *α* > 0 is a global parameter and *γ* > 0 is a subject-specific variable. *t*_0_ is a fixed age, here *t*_0_ = 70, so that *γ* is the progression score at 70 years. This model is partially motivated by Jedynak et al. [6, Fig. 4] and Bilgel et al. [7, Fig. 6, right]. In both applications, the progression score is an affine function of age for each subject, yet Eq. 1 provides a reasonable approximation. Solution of Eq. 1 is given by

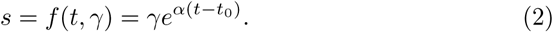

**Figure 4:**
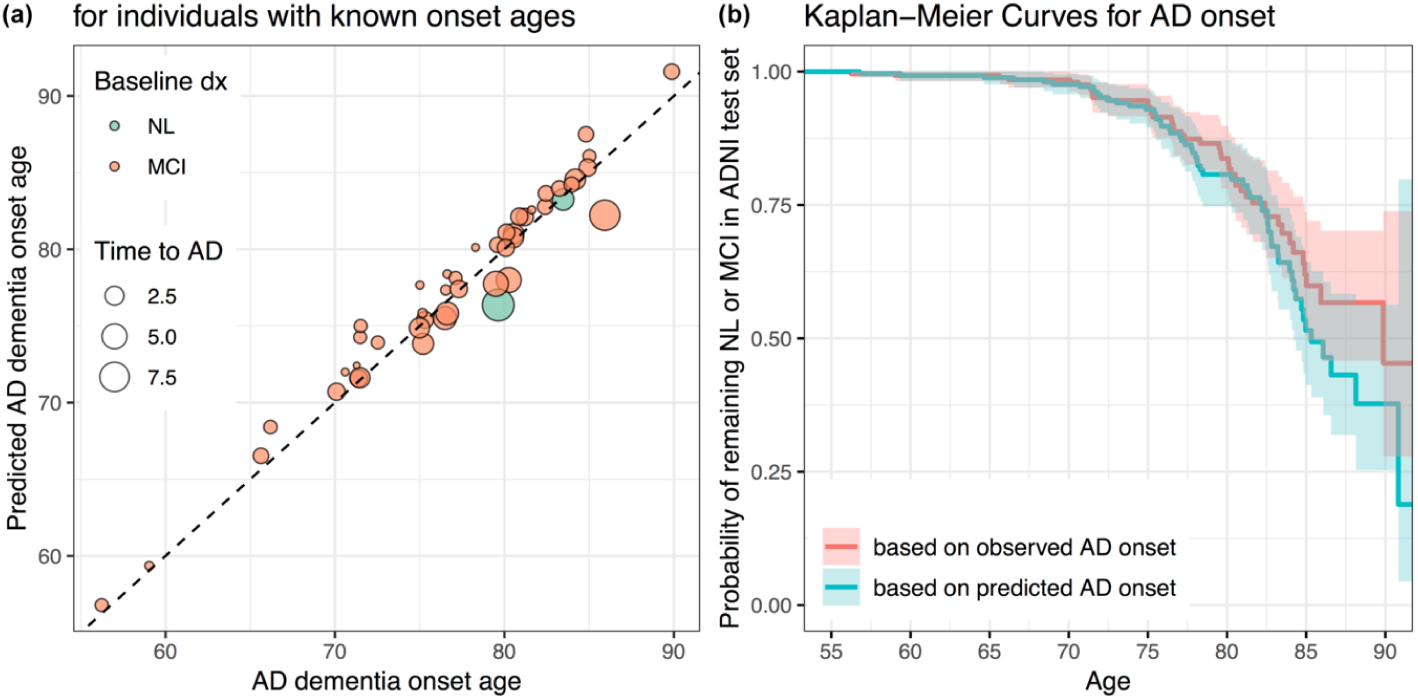
(a) AD dementia onset ages predicted using the linear regression model with PS + Age vs. observed AD dementia onset age (for individuals with known onset ages in the testing set). Time between baseline age and AD onset age (indicated by the size of the markers) varied between 0.48 and 9.0 years (median 1.6, IQR 1.0-3.0). (b) Kaplan-Meier curves based on observed (red) and predicted (blue) AD dementia onset ages.

Second, the vector of biomarker values at time *t* is modeled with a monotonic function of the progression score for each biomarker added to a correlated noise:

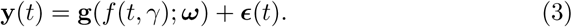

Monotonicity of **g** is required for model identifiability.

### Time from diagnosis

Diagnostic events include transitions from NL or MCI to AD. The model above allows for the computation of biomarker trajectories as function of time from a diagnosis. Let 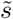 be the progression score corresponding to a diagnosis. For an individual with *s*(*t*_0_) = *γ*, the age 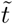 at diagnosis is, using Eq. 2,

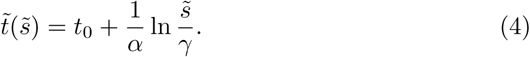

The time from diagnosis for this subject at age *t* is thus 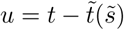. Biomarker trajectories as a function of time from diagnosis are characterized by the mapping

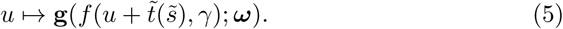

### Discrete-time model

We now describe the discrete-time model and the priors. Let *i* indicate a subject and *j* a visit. The subject-specific model is:

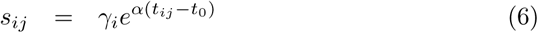

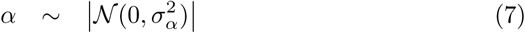

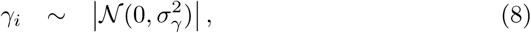

where *t_ij_* is the age of subject *i* at visit *j*, and |𝓝| is the half-normal distribution. We let to *t*_0_ = 70 and use half-normal hyperpriors on *σ_α_* and *σ_γ_*, with their scale parameters fixed at 0.05 and 5, respectively.

Given *K* biomarkers, their trajectory models parameterized by *ω* are given by:

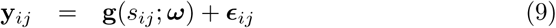

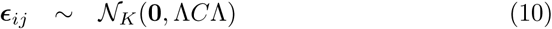

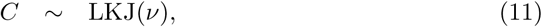

where Λ is a diagonal matrix with Λ_*kk*_ = *λ_k_*, *C* is a correlation matrix, and LKJ is the random correlation matrix distribution described by Lewandowski et al. [20]. We let *v* =1 for a uniform distribution over correlation matrices.

We consider logistic basis functions to characterize monotonic trajectories for each biomarker *k*:

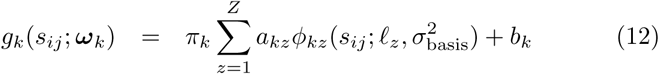

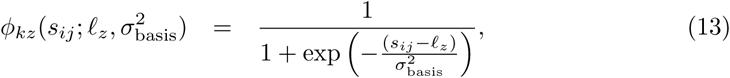

where *ω_k_* = {*π_k_, a_k1_,…, a_kZ_*, *b_k_*}, with the constraint that *a_kz_ >* 0 ∀*k,z* to ensure monotonicity.π_k_ is a categorical random variable with equally likely observations { — 1, +1} to determine whether the trajectory is decreasing or increasing, *ℓ* ∊ 𝕉^*Z*^ is a prespecified set of *Z* basis function locations, and σ^2^_basis_ determines the slant of each basis function.

Assuming that all biomarker observations *y_ijk_* have been standardized to have zero mean and unit standard deviation at baseline, we can let

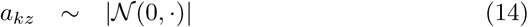

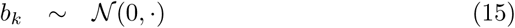

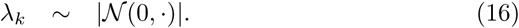

The standard deviation or scale parameter of each is fixed at a large value (i.e., 10) such that the priors are essentially uninformative. In the absence of data, these priors slightly favor flat “trajectories” for the biomarkers.

### Model fitting

We conducted Markov chain Monte Carlo sampling to generate posterior samples, using the No-U-Turn Sampler step method [21] for continuous variables, and a Metropolis-within-Gibbs step method optimized for binary variables for *π_k_*. Non-negative variables were log-transformed to allow for unconstrained optimization. For visits with missing biomarker measurements, only the available measurements contributed to the model fitting procedure. We used the PyMC3 Python package for model specification and fitting [22]. Model fitting was performed using training data.

We used *Z* = 5 logistic bases, with *ℓ_z_* equally spaced between 0 and 10 and σ^2^_basis_ = 1. We then fitted a series of models:

1. *σ_α_*, *σ_γ_*, and *C* were fixed at 0.05, 5.0, and the identity matrix *I_K×K_*, respectively. For tuning the sampling parameters, we obtained 300 samples, which were then discarded. The following 300 samples were used for model parameter estimation.
2. Next, we removed the constraint fixing the correlation matrix *C*. Samples obtained from the previous step were used to initialize, *ω_k_*, *λ_k_*, *α_i_*, *β_i_*, *π_k_*, and missing biomarker observations *y_ijk_*. We continued to fix *σ_α_* = 0.05 and *σ_γ_* = 5.0. This model was fitted using the longitudinal data set, with 200 tuning + 200 samples.
3. Finally, we removed the constraints fixing *σ_α_* and *σ_γ_*. Samples obtained from the previous step were used to initialize all previously estimated parameters, and we fitted this model using 2000 tuning + 2000 samples.

### Correlates of progression scores

To understand correlates of the individualized PS estimated by our model, we performed a multiple linear regression analysis using the training data set investigating age, education, sex, APOE *ɛ*4 status, and clinical diagnosis as predictors of PS at baseline.

### Biomarker trajectories as a function of time from AD dementia

For each of the last *M* = 200 Monte Carlo iterations, we drew a sample 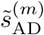 from the distribution of *s* at the first visit with an AD diagnosis among individuals who converted from NL or MCI to AD for the *m*^th^ Monte Carlo iteration (excluding individuals who reverted) and computed the curve 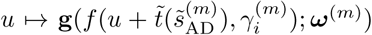 for each individual *i* in the training set. The resulting *MN* biomarker curves, where *N* is the number of individuals in the training set, were used to characterize the mean curve and its 95% interval. To check if our estimated trajectories agreed with the time from onset values available in the data set, we plotted the computed curves, superimposed on a scattergram of the biomarker data versus observed time from onset.

### Prediction of time to diagnosis

Age at the first occurrence of an AD dementia diagnosis for each individual who is NL or MCI at baseline was considered as the age at dementia onset. Using the known age at dementia onset data in the training set, we trained a linear regression model with time to dementia onset from baseline as the outcome and baseline progression score as the independent variable.

To make predictions for time to dementia onset in the testing set, we first used baseline age and baseline biomarkers to compute a progression score for each individual given the trained progression score model. The trained linear regression model was then applied to these baseline progression scores to make predictions for time to dementia onset for each individual in the testing set.

We assessed the performance of our onset age estimation by computing the root mean square error (RMSE) between the predicted and observed onset ages for individuals whose onset ages are known (i.e., for whom diagnostic conversion was observed in the longitudinal data set). This analysis is a biased reflection of the accuracy of onset age prediction given that it is restricted to those who convert to AD. To obtain measures based on all individuals regardless of their conversion status, we estimated Kaplan-Meier survival curves based on our predictions and observed onset ages. Survival curve estimation incorporates data for all individuals by assuming that AD conversion will occur after the last visit for individuals who did not convert during the study. For computing the KaplanMeier curves using observed data, event was defined as the first visit with an AD diagnosis following a visit with an NL or MCI diagnosis. We right-censored using age at last visit if the individual remained NL or MCI. For computing the Kaplan-Meier curves using predicted data, if onset was “predicted” to occur prior to baseline visit, age at onset was set to baseline age. We right-censored using age at last visit if the predicted onset age was greater than the age at last visit. We compared these two curves using the log-rank test *χ*^2^ statistic.

We repeated the linear regression models using age, each biomarker, age + each biomarker, all biomarkers, age + all biomarkers as independent variable(s). We also ran linear regression models using PS and its combinations with age and/or *γ* as independent variable(s). An intercept was included in all linear regression models. We then used permutation tests to compare the RMSE and *χ*^2^ of the best model without PS to those of the best model with PS. The permutation test involved randomly swapping the onset ages predicted by the models being compared and computing the differences in RMSE and *χ*^2^ between the two models. This was repeated 2000 times to obtain a distribution for RMSE difference and a distribution for *χ*^2^ difference under the null hypothesis of equality between models. The observed difference was then quantified against this null distribution using a two-tailed test.

### Reproducible research

We provide the code used for these analyses and all the software required to run it as a docker image accessible via https://hub.docker.com/r/bilgelm/bayesian-ps-adni/.

## 3. Results

Participant demographics are presented in Table 1. Difference in the proportions of NL and MCI diagnoses at baseline between training and testing sets was not statistically significant (Fisher’s exact test *p* = 0.79). There were no statistically significant differences between training and testing sets at baseline by diagnostic category in age, sex, race, education, APOE *ɛ*4 genotype, number of visits, or duration of follow-up.

### Fitted progression score model

Estimated biomarker trajectories as a function of PS are presented in Fig. 1. Resulting temporal QT revealed early changes verbal memory. The temporal evolution was also marked by changes in hippocampal volume and CSF A*β*_1-42_, tau and p-tau, as well as FDG-PET and remaining cognitive measures.

**Figure 1:**
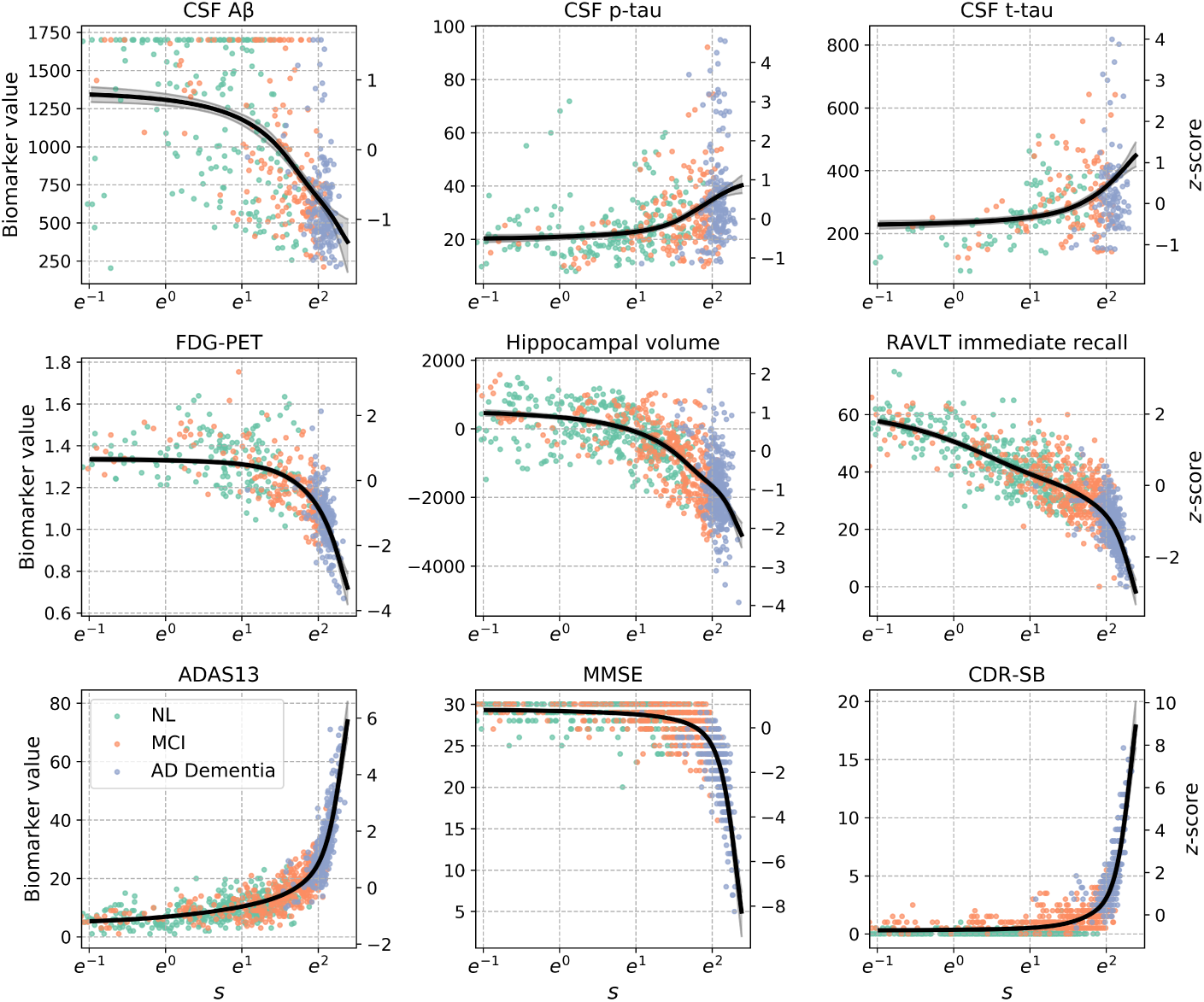
Estimated population-level biomarker trajectories as a function of progression score *s*. *s* is plotted on a natural logarithm scale so that the *x*-axis is linear in time. Mean trajectories are plotted in black, along with their 95% credible intervals. Longitudinal data points for 100 randomly sampled individuals per diagnostic category are shown. Biomarker *z*-scores, shown on the right-hand side *y*-axes, were computed using mean and standard deviations at baseline across 1369 in the training set.

### Correlates of progression scores

Box plots of the estimated subject-specific variable *γ* and baseline *s* revealed differences among cognitively normal, mild cognitively impaired, and demented individuals. These group-wise differences were greater than those observed with age (Fig. 2). Older age, male sex, fewer years of education, number of APOE *ɛ*4 alleles, and MCI or AD diagnoses were associated with higher PS at baseline (Table 2).

**Figure 2:**
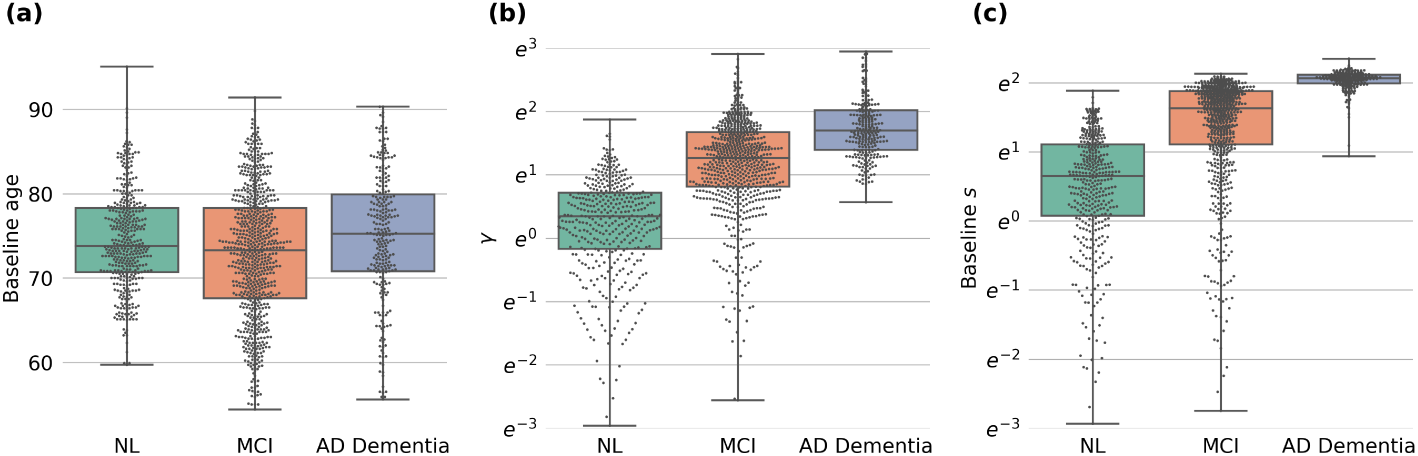
Box and swarm plots of (a) baseline age, (b) estimated *γ*, and (c) estimated progression score *s* at baseline by baseline diagnosis for individuals in the training set. *γ* and *s* are plotted on a natural logarithm scale. All pairwise comparisons were statistically significant (all *p* < 0.0013), with the exception of baseline age comparison between NL and AD.

**Table 2:** Result of multiple linear regression analysis investigating the associations of age, sex, education, APOE *ɛ*4 genotype, and clinical diagnosis with estimated progression score *s* at baseline. 5 individuals were excluded due to missing APOE information. 1364 observations were used to fit the model, yielding an *R^2^* = 0.635, an adjusted *R^2^* = 0.633, *F*-statistic= 336.7, Prob(*F*-statistic)< 0.0001. SE = Standard error.

| | Estimate | SE | $t$-statistic | $p$-value |
| --- | --- | --- | --- | --- |
| Intercept | -3.72 | 0.545 | -6.83 | < 0.001 |
| Age | 0.0869 | 0.006 | 14.1 | < 0.001 |
| Male sex | 0.303 | 0.089 | 3.40 | 0.001 |
| Education | -0.0619 | 0.016 | -3.98 | < 0.001 |
| APOE $\epsilon 4$ heterozygous | 0.719 | 0.094 | 7.63 | < 0.001 |
| APOE $\epsilon 4$ homozygous | 1.34 | 0.158 | 8.50 | < 0.001 |
| MCI | 2.45 | 0.103 | 23.9 | < 0.001 |
| AD dementia | 5.13 | 0.132 | 38.8 | < 0.001 |

### Time from diagnosis

Biomarker trajectories as a function of time from AD diagnosis are shown in Fig. 3.

**Figure 3:**
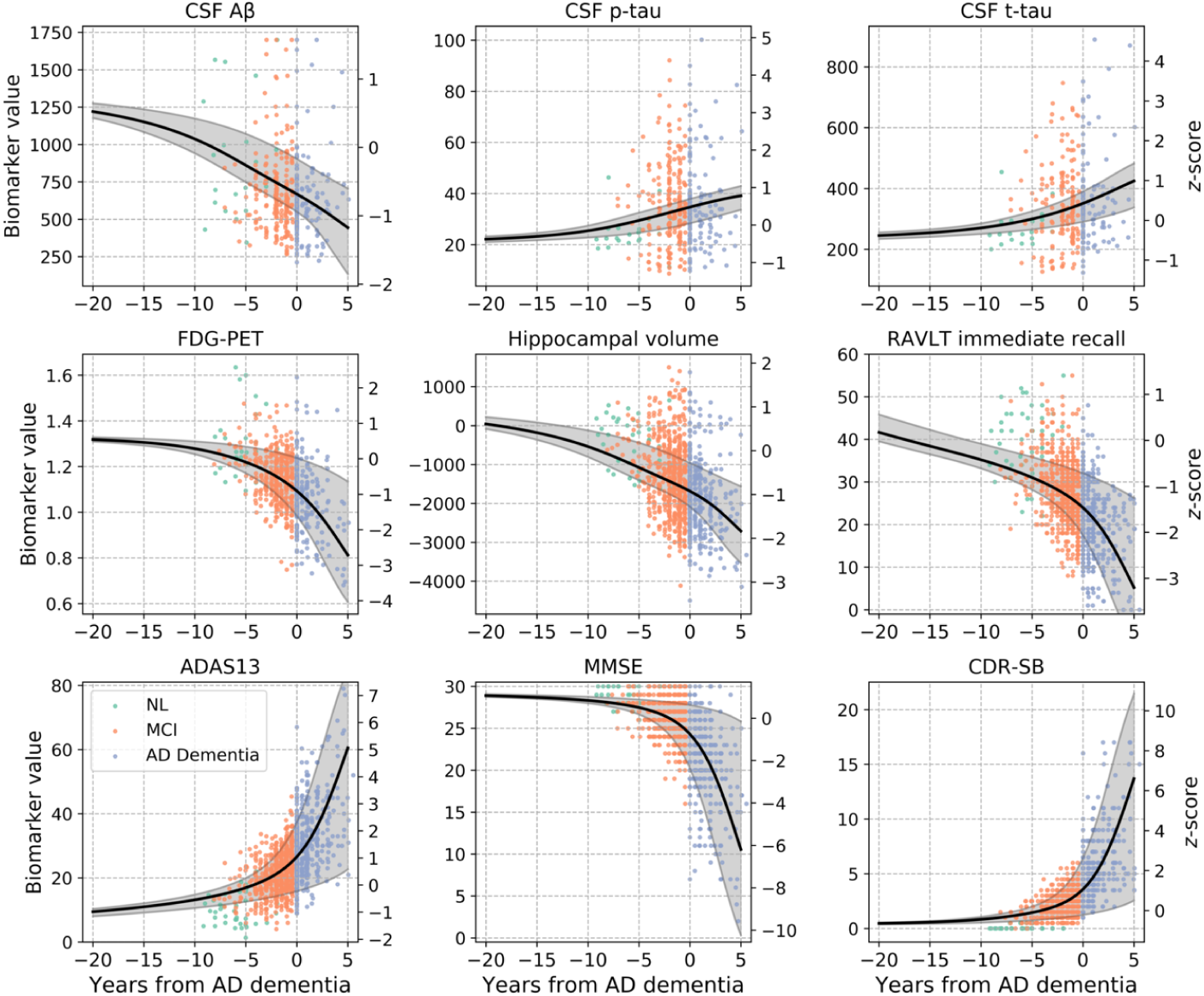
Estimated biomarker trajectories as a function of time from initial AD dementia diagnosis are shown with the black curves, and the shaded areas depict 95% credible intervals. Biomarker *z*-scores, shown on the right-hand side *y*-axes, were computed using mean and standard deviations at baseline across 1369 individuals in the training set. Note that observed time from AD was not used in the estimation of the biomarker trajectories shown; trajectories were obtained using the model fit. Scattergrams of observations are shown as a function of observed time from AD, color-coded by diagnosis, to allow for a visual assessment of the agreement of the estimated trajectories with underlying data.

Among prediction models that did not include PS as an independent variable, the model with the lowest *χ^2^* was the one with all 9 biomarkers (*χ^2^* = 20.5, RMSE = 1.58), whereas additionally including age yielded the model with the lowest RMSE (*χ^2^* = 21.1,RMSE = 1.57) (Table 3). Among models with PS, including age in addition to PS yielded the model with the lowest *χ^2^* (*χ^2^* = 1.68,RMSE = 1.49), whereas the model with PS only had the lowest RMSE (*χ^2^* = 2.00,RMSE = 1.48).

**Table 3:** Prediction results based on linear regression models, presented for a select subsample of investigated models. RMSE = root mean square error.; Max. abs. err. = maximum absolute error; *χ^**2**^* = log-rank test statistic; Max. vert. dist. = maximum vertical distance between the survival curves based on predicted onset and observed onset; RAVLT imm. = Rey Auditory Verbal Learning Test immediate recall (sum across learning trials); PS = progression score.

| Independent variable(s) | Measures based on<br>those who convert |  | Survival curve<br>based measures |  |
| --- | --- | --- | --- | --- |
| | RMSE | Max.<br>abs. err. | $\chi^2$ | Max.<br>vert. dist. |
| Age | 1.80 | 6.78 | 46.6 | 0.491 |
| RAVLT imm. | 1.64 | 5.52 | 35.7 | 0.438 |
| RAVLT imm. + Age | 1.64 | 5.46 | 34.5 | 0.437 |
| All biomarkers | 1.58 | 4.67 | 20.5 | **0.428** |
| All biomarkers + Age | 1.57 | 4.63 | 21.1 | 0.437 |
| PS | **1.48** | 3.79 | 2.00 | 0.453 |
| PS + $\gamma$ | **1.48** | 3.75 | 1.72 | 0.453 |
| PS + Age | 1.49 | **3.70** | **1.68** | 0.453 |

Results of age at AD onset prediction based on the linear regression model with PS and age are shown in Fig. 4. The Kaplan-Meier curve computed using predicted onset ages tracked the observed curve closely (Fig. 4b). We used the log-rank test to compare the curve based on predicted onset ages to the curve based on observed onset ages. The difference between the curves based on predicted and observed onset ages was not statistically significant based (log-rank test *p* = 0.2).

We compared the performance of the model with all 9 biomarkers to that of the model with PS + Age, since these models had the lowest *χ^2^* measures. Based on permutation testing, the difference in RMSE (*p* = 0.75) or *χ^2^* (*p* = 0.08) of the two models was not statistically significant. Unlike the predictions based on PS + Age, predictions based on all 9 biomarkers yielded a survival curve that was different from the survival curve based on the observed ages (log-rank test *p* < 0.0001).

## 4. Discussion

We presented a model for aligning short-term longitudinal data across individuals to characterize long-term trajectories of a collection of biological and cognitive measurements that are implicated in AD. We applied our model to individuals who are cognitively normal, have MCI or AD dementia to estimate a QT of AD progression using the ADNI dataset and recommendations presented in the Alzheimer’s Disease Modelling Challenge. We demonstrated that progression along our estimated temporal QT reflects AD processes by showing its association with clinical diagnosis and known AD risk factors, as well as by quantifying the accuracy of our model in predicting age at onset of AD dementia. The ability of our model to leverage short-term data to obtain long-term biomarker trajectories is highly relevant to the current need in clinical trial design for predicting who will develop cognitive impairment and AD dementia over time.

We found that the PS computed based on AD-related neuropathology, neurodegeneration, and cognitive measures were associated with age, APOE ε4 positivity, which is the most influential known genetic risk factor for sporadic AD [23], and clinical diagnoses of MCI and AD. We also observed higher PS among men and a negative association between PS and years of education. Associations with APOE *ɛ*4 and clinical diagnoses provided evidence that our QT, which was constructed without these pieces of information, is reflective of AD progression. While we recognize that there is heterogeneity in individual presentations across the biomarkers considered in our model and that not every individual included in our analyses is necessarily on an AD pathway, the strong associations we observed between PS and AD diagnosis as well as AD risk factors suggest that the estimated trajectories are mainly indicative of age-and disease-related changes that occur along the progression towards AD dementia.

Our QT was consistent with our previous findings in ADNI [6]. By modifying our previously described Progression Score Models [6, 7] to include a global parameter governing the relationship between disease stage and rate of progression, we were able to depict biomarker trajectories as a function of time from AD diagnosis. This innovation enabled a more easily interpretable characterization of the natural history of AD starting in preclinical stages.

Our method achieved < 1.5-year RMSE in predicting age at onset of AD over the course of 0.48 to 9.0 years. It is difficult to compare the performance of our prediction with previous methods given differences in samples and model features used. Oxtoby et al. [5] reported an RMSE of 1.3 years for predicting age at onset over the course of 3.0 years in the context of dominantly-inherited Alzheimer’s disease based on CSF, MRI, and positron emission tomography biomarkers. Vogel et al. [24] reported an *R^2^* value of 0.15 using the ADNI data set for predicting years to conversion to MCI or AD based on functional and structural MRI measures. It is necessary to continue model development in this area to enable more accurate onset age predictions over longer time intervals.

A limitation of our model is the simplifying assumption of a single pathway of biomarker changes from a cognitively normal state to AD dementia. It is known that there is heterogeneity in longitudinal progression in AD. Studying individuals who deviate from the estimated QT may be informative in understanding this heterogeneity, and models that relax the assumption of a single progression pathway will be necessary for a detailed analysis. We observed the greatest variability around trajectory estimates for CSF A*β*_1-42_. This suggests that the timing of CSF A*β*_1-42_ can vary at the individual level in relation to changes in other biomarkers included in this study. Another limitation of our study is that higher PS indicates both age-and AD-related changes in the examined biomarkers. Understanding changes that occur with AD as distinct from age-related changes is clinically important. However, this is a challenging goal given that there is no scientific consensus regarding the definition of healthy aging. Several studies have approached this problem by including age and other demographic variables as covariates in addition to a disease stage variable [11, 25, 26] or by regressing out their effects prior to model fitting [5, 27, 28], whereas others have not included covariates in order to characterize trajectories that explain the natural history of AD dementia, including biological and cognitive changes that occur due to aging in addition to AD [4, 6–9, 13, 15]. We formulated our goal with this study as the characterization of the natural history of AD dementia, including biological and cognitive changes that occur due to aging in addition to AD pathology, and therefore did not include an adjustment for age or any other demographic variables.

In conclusion, our method allows for the estimation of individualized latent disease progression indicators and population-level biomarker trajectories from longitudinal data. The estimated temporal QT of AD provides a mechanism for localizing individuals along biomarker trajectories based on multivariate data. Individualized progression scores can be used longitudinal composites to investigate associations with risk factors and outcomes. Furthermore, the ability to obtain individualized estimates of age at AD onset can allow for better participant recruitment in clinical trials aimed at preclinical AD.

## Acknowledgments

Data collection and sharing for this project was funded by the Alzheimer’s Disease Neuroimaging Initiative (ADNI) (National Institutes of Health Grant U01 AG024904) and DOD ADNI (Department of Defense award number W81XWH-12-2-0012). ADNI is funded by the National Institute on Aging, the National Institute of Biomedical Imaging and Bioengineering, and through generous contributions from the following: AbbVie, Alzheimer’s Association; Alzheimer’s Drug Discovery Foundation; Araclon Biotech; BioClinica, Inc.; Biogen; Bristol-Myers Squibb Company; CereSpir, Inc.; Cogstate; Eisai Inc.; Elan Pharmaceuticals, Inc.; Eli Lilly and Company; EuroImmun; F. Hoffmann-La Roche Ltd and its affiliated company Genentech, Inc.; Fujirebio; GE Healthcare; IXICO Ltd.; Janssen Alzheimer Immunotherapy Research & Development, LLC.; Johnson & Johnson Pharmaceutical Research & Development LLC.; Lumosity; Lundbeck; Merck & Co., Inc.; Meso Scale Diagnostics, LLC.; NeuroRx Research; Neurotrack Technologies; Novartis Pharmaceuticals Corporation; Pfizer Inc.; Piramal Imaging; Servier; Takeda Pharmaceutical Company; and Transition Therapeutics. The Canadian Institutes of Health Research is providing funds to support ADNI clinical sites in Canada. Private sector contributions are facilitated by the Foundation for the National Institutes of Health (www.fnih.org). The grantee organization is the Northern California Institute for Research and Education, and the study is coordinated by the Alzheimer’s Therapeutic Research Institute at the University of Southern California. ADNI data are disseminated by the Laboratory for Neuro Imaging at the University of Southern California.

This research was supported in part by the Intramural Research Program of the National Institute on Aging, National Institutes of Health as well as the Portland Institute for Computational Science and its resources acquired using NSF Grant DMS 1624776.

## Footnotes

1 Data used in preparation of this article were obtained from the Alzheimer’s Disease Neuroimaging Initiative (ADNI) database (adni.loni.usc.edu). As such, the investigators within the ADNI contributed to the design and implementation of ADNI and/or provided data but did not participate in analysis or writing of this report. A complete listing of ADNI investigators can be found at: http://adni.loni.usc.edu/wp-content/uploads/how_to_apply/ADNI_Acknowledgement_List.pdf

